# Early-Career Coordinated Distributed Experiments: Empowerment Through Collaboration

**DOI:** 10.1101/704502

**Authors:** Ada Pastor, Elena Hernández-del Amo, Pau Giménez-Grau, Mireia Fillol, Olatz Pereda, Lorea Flores, Isis Sanpera-Calbet, Andrea G. Bravo, Eduardo J. Martín, Sílvia Poblador, Maite Arroita, Rubén Rasines-Ladero, Celia Ruiz, Rubén del Campo, Meritxell Abril, Marta Reyes, Joan Pere Casas-Ruiz, Diego Fernández, Núria de Castro-Català, Irene Tornero, Carlos Palacin-Lizarbe, María Isabel Arce, Juanita Mora-Gómez, Lluís Gómez-Gener, Silvia Monroy, Anna Freixa, Anna Lupon, Alexia María González-Ferreras, Edurne Estévez, Pablo Rodríguez-Lozano, Libe Solagaistua, Tamara Rodríguez-Castillo, Ibon Aristi, Aingeru Martínez, Núria Catalán

## Abstract

Coordinated distributed experiments (CDEs) enable the study of large-scale ecological patterns in geographically dispersed areas, while simultaneously providing broad academic and personal benefits for the participants. However, the effective involvement of early-career researchers (ECRs) presents major challenges. Here, we analyze the benefits and challenges of the first CDE exclusively led and conducted by ECRs (i.e. ECR-CDE), which sets a baseline for similar CDEs, and we provide recommendations for successful CDE execution. ECR-CDEs achieve most of the outcomes identified in conventional CDEs as well as extensive benefits for the young cohort of researchers, including: (i) receiving scientific credit, (ii) peer-training in new concepts and methods, (iii) developing leadership and communication skills, (iv) promoting a peer network among ECRs, and (v) building on individual engagement and independence. We also discuss the challenges of ECR-CDEs, which are mainly derived from the lack of independence and instability of the participants, and we suggest mechanisms to address them, such as resource re-allocation and communication strategies.

## 1 Introduction

Understanding global environmental questions requires the ability to generalize the results found at one site to other environments. Coordinated distributed experiments (CDEs), i.e. manipulative experiments run in parallel by geographically dispersed teams, constitute a powerful tool to test ecological questions across large geographical scales. CDEs are characterized by the synchronous use of low-cost standardized sampling methodologies and represent the state-of-the-art in collaborative science (Fraser et al., 2013; Borer et al., 2014). In ecological research, successful CDEs have led to key advances in our understanding of processes such as litter decomposition (RivFunction project; Woodward et al., 2012; LIDET Gholz et al. 2000) or nutrient dynamics (LINX projects; LINX collaborators 2014). Moreover, CDEs offer benefits for participants that are not typically measured, such as the development of scientific networks, training in new methodologies, and personal satisfaction (Goring et al., 2014; LINX collaborators, 2014).

The participation of early-career researchers (ECRs) in CDEs is key to ensure innovative and fruitful science (Callaway, 2015). The ECRs might also benefit individually from involvement in a CDE network, for example, by increasing their academic visibility within the scientific community. However, ECRs face greater challenges than their senior colleagues (Goring et al., 2014). First, ECRs are usually reliant on the participation of their group leader and associated economic resources. Second, the short-term positions ECRs usually hold might hamper effective ECR involvement in CDEs and constrain the benefits of their participation. Finally, credit for their work within a CDE might be diluted in favor of more reputable scientists (Merton, 1968). Thus, it is crucial to find appropriate tools for ECRs to take full advantage of participating in CDEs and constructively build on a long-term positive scientific culture.

Recently, some scientific societies have promoted the collaboration of young researchers by funding CDEs exclusively targeted to ECRs (hereafter referred to as ECR-CDEs). Through specific funding calls, these societies support projects that have a CDE structure and both calls and projects are open to all the ECRs within a society. ECR-CDE projects have the potential for ecological breakthroughs and to positively impact ECR careers. However, because of the novelty of these instruments, these benefits remain undetermined. Here, we use our collective experience as participants in the first ECR-CDE (2013-2015; see Supplementary material) to (1) discuss the benefits and challenges of ECR-CDEs and (2) provide recommendations for the successful development of future initiatives. Overall, we endorse ECR-CDEs as effective tools to assess large-scale ecological questions while empowering ECRs.

## 2 Benefits of collaborating through ECR-CDEs

Participating in ECR-CDEs extends the benefits of common CDEs for young researchers across academic, training, and personal development areas (Fig. 1). To evaluate the participant benefits of the first ECR-CDE, an anonymous online survey was launched at the end of the project. The survey collected information on career stage (*see* supplementary Fig. 1 for categories), research field, age, time devoted to the project, opinion about the project management and development and suggestions for improvement. The results of the survey (Fig. 2) are discussed here onwards within the context of a broad evaluation of potential benefits of ECR-CDEs.

**Figure 1.**
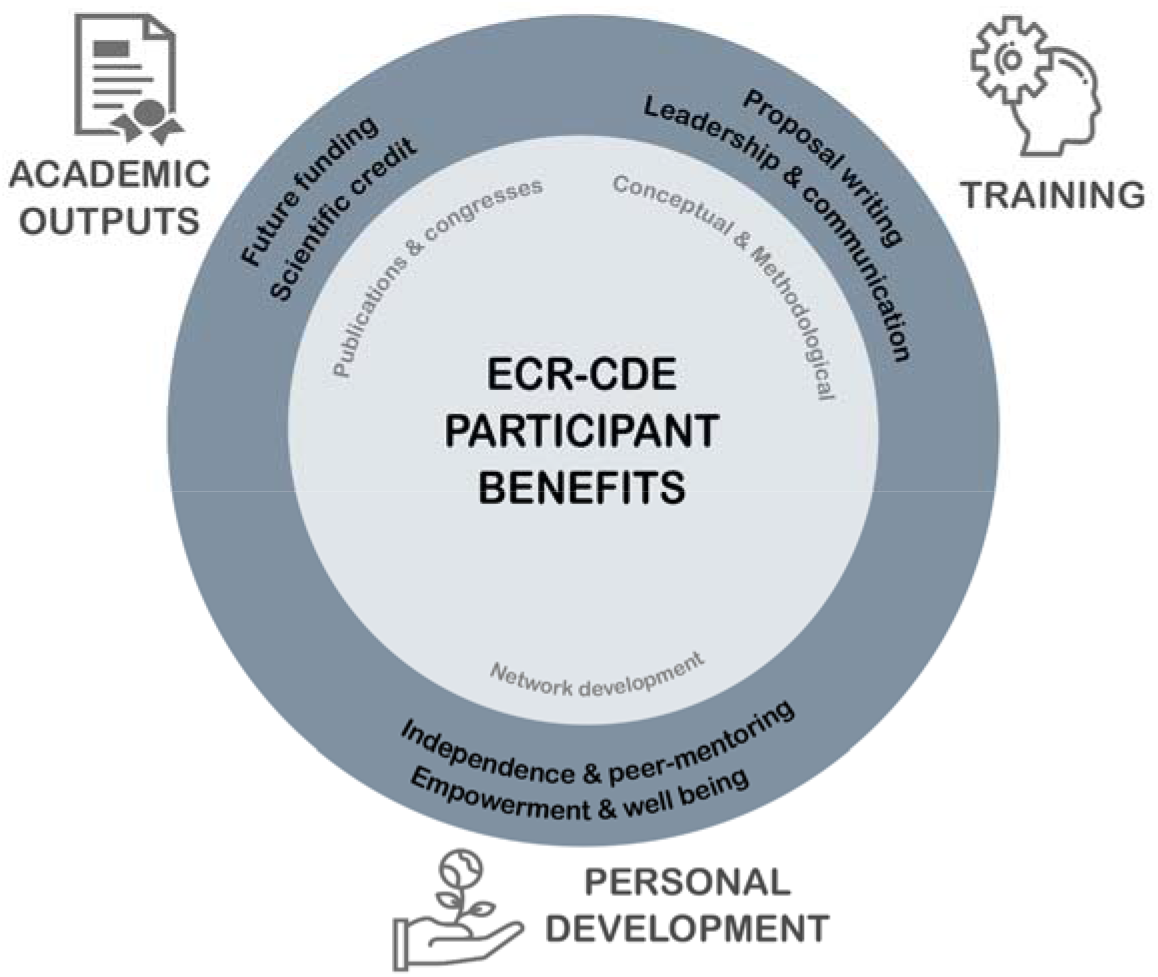
Benefits for participants identified for CDE (inner light circle; Goring et al. 2014) and extended benefits for ECR-CDE (outer circle; identified in the present study) in three main areas: academic outputs, training and personal development.

**Figure 2.**
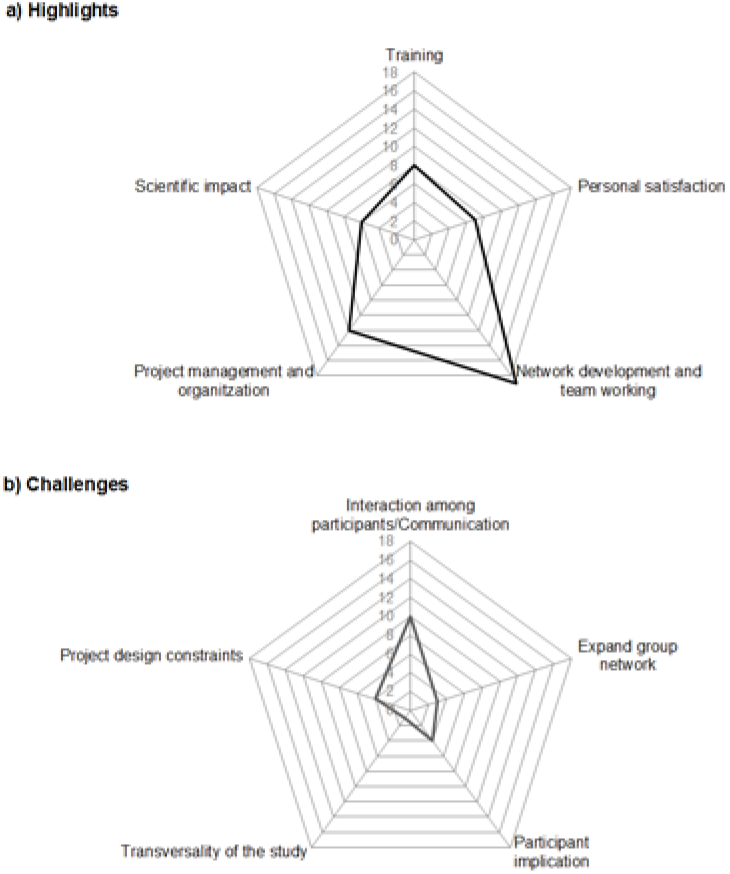
Results of the satisfaction survey were ECRs determined the (a) highlights and (b) challenges of the pioneer ECR-CDE. The values correspond to the number participants who cited the different aspects of the project in an open question of the anonymous survey (note that not all the participants identified challenges).

### 2.1 Academic impact

The CDE structure allows addressing complex environmental questions on large geographic scales (Fraser et al., 2013), thus CDEs may produce high-impact *publications and communications*. However, one of the most critical challenges for ECRs in collaborative environments is achieving individual recognition (Goring et al. 2014), which is often awarded to the most senior participants (Merton, 1968). ECR-CDEs grant the *scientific credit* to the ECR cohort, promoting their visibility, and thus increasing the likelihood of successful *future funding applications* (Fig. 1).

### 2.2 Training

ECRs also benefit from participation in a collaborative project by gaining knowledge and skills beyond those acquired during the student stage (Goring et al., 2014; Fig. 1). Indeed, *training on new concepts and methodologies* was a highly valued outcome by the ECRs (Fig. 2a). In contrast to regular CDEs, methodological training in ECR-CDEs is organized as a dispersed learning structure, in which students build their own learning network, a proven pedagogical tool (Boud & Lee, 2005. The shared effort devoted to obtaining comparable data is an effective implementation of action-oriented learning. In addition to training on new concepts and methodologies, which is common to conventional CDEs, projects exclusively targeting ECRs provide a great opportunity to submit the first research proposal of their careers in a fair competitive league. *Training on proposal writing* is key for successful career development (Porter 2004; Davis 2009), and evaluation of the proposals by a specialized international committee is a great opportunity to gain insight for future project calls. Moreover, the coordination of a CDE provides unparalleled *training on leadership skills and project management* for the project coordinators (Fig. 1). These abilities are required for the most prestigious scientific grants (e.g. the Starting Grant of the European Research Council) and are an asset for successful scientific career development (Leiserson & McVinney, 2015).

Participation in an ECR-CDE promotes training in *skills and strategies on how to collaborate, communicate, and share effectively* (Fig. 1). The development of the ECR-CDE required strong communication practices during all the steps of the project (Supplementary Figure 2). Although explicit communication training is rare in graduate programs (Cheruvelil et al., 2014), scientists require a broad range of communication skills in order to engage with other researchers and decision makers in collaborative processes (*see* translational ecology; Schwartz et al., 2017). Moreover, communication among peers enables effective collaboration, crucial for high-performing science (Bercovitz & Feldman, 2011; Lortie et al., 2012). ECR-CDEs constitute valuable experience for this end.

### 2.3 Personal development

ECR-CDEs can significantly contribute to the personal development of ECRs. While the reasons to step aside from a research career are multidimensional and may include personal and institutional aspects, the feeling of isolation and the lack of adequate socialization have been highlighted as central motives to discontinue an academic career (Pyhältö & Keskinen, 2012; Litalien & Guay, 2015; Castelló et al., 2017). Accordingly, *network development* was the most valued benefit of the pioneer ECR-CDE (Fig. 2a). By developing their own network of collaborators and participating in a project apart from their supervisor, ECRs gain autonomy and acquire a wider scientific perspective, becoming *empowered* and more *independent researchers* (Fig. 1). Moreover, having a network of collaborators *promotes peer-mentoring*, accelerating the transition from apprentice to colleague, thus helping ECRs to advance in their scientific careers (Laudel & Gläser, 2008). Peer collaboration throughout ECR-CDEs may also have positive consequences on the *socio-psychological well-being* of the participants (Fig. 1). Feelings of empowerment and social bonding through participation in an ECR-CDE may promote research engagement and reduce stress, exhaustion, and anxiety (Stubb et al., 2011), mental health issues that are common among graduate students (Evans et al., 2018). The benefits of ECR-CDEs could have greater effects on and therefore potentially decrease drop-out ratios of particularly vulnerable ECR groups, such as young women, (Shaw & Stanton, 2012; Sánchez-Montoya et al., 2016). Although it is still early as most participants have only transitioned from PhD students into post-docs, in the inaugural ECR-CDE, the drop-out ratio of women was only 5 % (n=2) and none of the initial post-docs have changed their career paths.

## 3 Challenges, actions and opportunities

The exclusive participation of ECRs confers ECR-CDEs some particularities that can negatively affect their development. Here, we identify and analyze these challenges and suggest strategies for the effective participation of young scientists in other CDEs (Table 1). We further discuss opportunities for these ECR-CDE initiatives.

**Table 1.** Particularities of the participants’ challenges, drawbacks and recommendations for CDEs with exclusive participation of early-career researchers (ECR-CDE) compared with a conventional CDE

| Particularities of participants | Potential drawbacks | Recommended actions |
| --- | --- | --- |
| *Lack of independence at economic and professional levels:* |  |  |
| - *Economic* | The groups lack their own funding resources | - Project core funding has to be distributed to groups<br>- Participants need to secure additional funding<br>- Budget allocation focused on collaboration and inclusiveness to the project (e.g. meetings attendance) rather than in experimental supplies |
| - *Professional (i.e. the participants have to "ask for permission" from their supervisors)* | Decreases the number of participants and their level of involvement | - Development and change of reward measures for participants (and supervisors)<br>- Consider ancillary projects that could be used for participants individual projects (e.g. MSc or PhD thesis) |
| *Lack of professional stability* | Increases CDE outcome uncertainty | - Reduce the duration of the CDE<br>- Assure the participant receives the project benefits |
| *Exclusive ECR participation* | Lack of intergenerational exchange | - Encourage individual exchange with supervisors<br>- Creation of a mentoring committee by the funding organization.<br>- Integration of the CDE within a scientific society or community |

First, the *autonomy of ECRs* is often limited at both economic and professional levels (Table 1). Collaboration is often expensive for the project, as a significant part of the resources need to be devoted to coordination, and for the participants as they might need to cover part of the sampling or meeting costs (Cummings & Kiesler, 2007). These costs particularly affect young scientists, who frequently lack discretionary funds. To address economic limitation, careful management of the available CDE funding is crucial. We suggest that in ECR-CDEs, particularly when the budget is limited, project management and the organization of in person workshops should be prioritized. Meetings promote participant involvement in the project, but also increase cost-effectiveness by facilitating equipment sharing, finding alternative facilities for analyses, or modifying the proposed methods. While discussions are the basis of scientific development, material provision can be achieved through resource re-allocation (Bollen et al. 2017). First, the *autonomy of ECRs* is often limited at both economic and professional levels (Table 1).

Collaboration is often expensive for the project, as a significant part of the resources need to be devoted to coordination, and for the participants as they might need to cover part of the sampling or meeting costs (Cummings & Kiesler, 2007). These costs particularly affect young scientists, who frequently lack discretionary funds. To address economic limitation, careful management of the available CDE funding is crucial. We suggest that in ECR-CDEs, particularly when the budget is limited, project management and the organization of in person workshops should be prioritized. Meetings promote participant involvement in the project, but also increase cost-effectiveness by facilitating equipment sharing, finding alternative facilities for analyses, or modifying the proposed methods. While discussions are the basis of scientific development, material provision can be achieved through resource re-allocation (Bollen et al. 2017). Moreover, focused development and discussions during in person events expedite reaching project milestones. Accordingly, participants in the first ECR-CDE recommended maximizing team interactions (Fig. 2b). Indeed, group productivity is strongly associated with in person meetings (Hampton & Parker, 2011). Among other activities, teambuilding exercises should also be encouraged during these meetings to develop interpersonal skills (Cheruvelil et al., 2014).

Another difficulty to address is the *limitation of autonomy at a professional level*, as ECR activities are often contingent on supervisor approval (Goring et al., 2014; Sala-Bubaré & Castelló, 2016). ECR-CDE benefits (Fig. 1) are usually not evaluated by the most common metrics to measure excellence in science and do not directly impact the supervisors’ scientific record. As an unfortunate consequence, supervisors might not consider ECR participation in these projects relevant to professional development and might disregard CDE project calls. Indeed, existing measures of reward may not be suitable to encourage collaboration, especially among ECRs (Goring et al., 2014). Criticism of metric-based success is rising (Fischer et al. 2012) and the need for a paradigm shift in the evaluation of scientific performance is becoming more apparent (Abbott et al., 2010; Dinsmore, 2014). An alternative is to include extended measures of success that assess non-tangible and long-term benefits such as personal satisfaction or group development (Goring et al., 2014). Extended metrics based on participants’ future success (Acuna et al. 2012), received training, team building, network development and open science (Wilsdon et al. 2017), should provide further incentives for supervisors to support ECR participation in ECR-CDEs.

The *lack of stability* of young scientists must also be considered (Table 1). Although long-time project windows are usually needed for the development of effective cooperative structures, these long-time frameworks hampers the inclusion of young scientists in the research outputs (Goring et al., 2014). Long project durations might result in detachment of the participants, thus the restriction of the experimental phase of the project to one or two years might better fit the timelines of ECRs, decreasing outcome uncertainty. It is also crucial that the project coordinators keep track of the participants, despite changes in their affiliations or career status, to make sure they receive the benefits of their initial participation.

Third, although similar age and position in career stage creates close-knit networks (Freeman & Huang, 2014), the exclusive inclusion of a unique scientific-stage cohort could limit the interaction with senior researchers and other career stages (Table 1). We suggest regular discussions with the participants' advisors and the appointment of a small committee of independent senior mentors. For example, this committee could be made of the same scientists that evaluated the projects during the call. Additionally, ecological or scientific societies in general can provide excellent mentoring networks that could support the ECR-CDEs to achieve the milestones of the projects while ensuring intergenerational exchange. While ECR-CDEs are not the only valid approach to facilitate ECRs’ career development, specifically not to the detriment of conventional CDEs or supervisor-student schemes, the development of novel initiatives might improve traditional mechanisms and positively impact other spheres related to academic culture for ECRs.

In the field of ecology, around sixty scientific societies are currently active in Europe. Initiatives like the one presented here could target thousands of ECRs in a specific ecological field in Europe alone. The second Iberian Association of Limnology ECR-CDE is currently running, with the participation of more than 60 young researchers, and the third is already in preparation. The achievements of this initial experience stimulated the European Federation for Freshwater Sciences to launch analogous calls for European ECRs in 2016 and 2018. The first of those united a team of 47 scientists from 25 institutions across 11 countries (https://freshproject-eurorun.jimdo.com/). The second, includes more than 100 scientists in 29 teams across 12 countries (https://freshproject-urbanalgae.jimdo.com/). We recommend that ECR-CDE initiatives should be open to participation for all interested ECRs, thus not restrict the positive effects to an exclusive group. Moreover, the first ECR-CDE can act as a template to apply manipulative-experiment approaches across other collaborative initiatives among non-specialized participants, such as in citizen science (Cohn 2008). Those collaborative initiatives should be encouraged to support the integration of transnational ecological research (Hoekman, Frenken, & Tijssen, 2010).

## 4 Concluding remarks

ECR-CDEs can advance the development of ecological and environmental research, while simultaneously advancing the careers of ECRs across disciplines (e.g. through training, networking and empowerment). This study shows the potential of ECR-CDEs as tools to 1) develop a scientific community based on values as altruism, willpower, and clear communication rather than on ego and individualism; and 2) foster the development of fruitful and collaborative science. The insights gained from the first ECR-CDE are linked to aspects of scientific culture that have strong impacts on career development and that should help to build a more socially sensitive culture in ecological sciences.

## 6 Author Contributions

AP and NC acquired funding, administered the project, conceptualized and drafted the manuscript. All authors contributed to review and editing this final version

## 8 Acknowledgments

We thank the support of all the affiliated institutions for the accessibility to laboratory premises, especially the Limnology Department of Uppsala University and the Department d’Ecologia of the Universitat de Barcelona. We greatly thank all the participants of DOMIPEX project. We are thankful to A. Sala-Bubaré for her helpful comments on an early version of this manuscript. We are deeply grateful to Anne M. Kellerman for the English corrections and comments on the manuscript.

