## Supplementary material for "Early-Career Coordinated Distributed Experiments: Empowerment Through Collaboration"

### 1 Supplementary Text. Structure and management of the first ECR-CDE

The studied ECR-CDE was selected from a competitive call supported by the Iberian Association of Limnology (AIL) and evaluated by a specialized international committee. The call aimed to tackle projects proposing spatial-temporal frameworks in freshwater sciences that could not be covered by a single group of researchers, so that a collaborative approach was needed. The project aimed to study metabolism and biogeochemical uptake among streams spanning a wide range of characteristics. It was opened for participation to the whole cohort of ECRs affiliated to the Iberian Association of Limnology (i.e. ≈ 150 members in 2014). The final project team included 42 ECRs from 15 institutions (in 2015) from Switzerland, Germany, Sweden and Spain. The age of the participants ranged from 23 to 37 years old and most of them were PhD students (67%) and short-term Post-Docs (21%; Supplementary Figure 1). Two thirds of the participants in the project were women, in agreement with their higher proportion in early career stages (Sánchez-Montoya et al. 2016).

The CDE had a reduced budget (4500 €; VAT included) and operated with a disperse organization (sensu Goring et al., 2014), whereby participant groups were self-organized, geographically distributed, and communicated regularly with the project coordinators. Communication tools included online meetings, emails and a Blog (http://jiail.blogspot.com.es/) frequently updated for communication with the participants and the public. The participant groups conducted two field sampling campaigns in eleven streams (Pastor et al. 2017; Catalán et al. 2018). Site selection and team creation was a long feedback process implying strong communication efforts (Supplementary Figure 2), although the previous knowledge about the sites of the team members strongly facilitated the selection and compilation of background information. A face-to-face kick-off meeting was held for team consolidation (July, 2014), where the initial protocol draft was explained, discussed and improved to finally be integrated in a detailed protocol prepared by the project coordinators (DOI: 10.13140/RG.2.2.14144.74245). Data treatment and processing was based on the participant's expertise, such as the application of nutrient uptake and metabolism metrics. This iterative communication was maintained throughout the project stages and defined the field, laboratory, and data treatment methods (Supplementary Figure 2).

### 2 Supplementary Figures and Tables

#### 2.1 Supplementary Figures

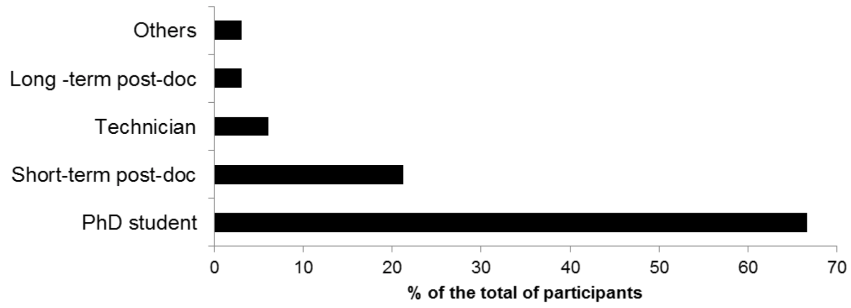


**Supplementary Figure 1.** Percentage of participants by career category at the beginning of the ECR-CDE (in 2013)

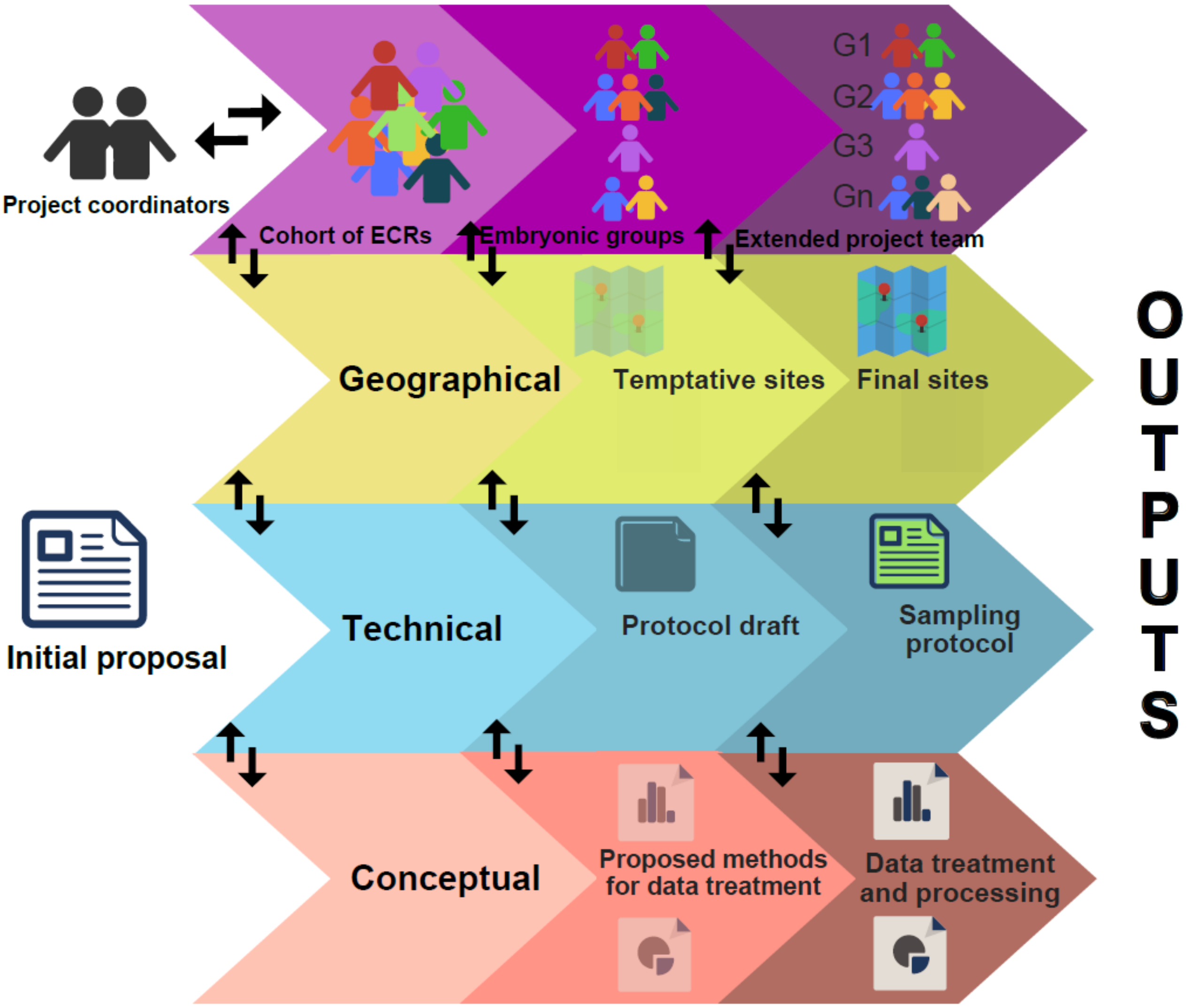


**Supplementary Figure 2.** ECR-CDE development. The geographical, technical and conceptual facets were developed from the initial proposal through an iterative communication process in parallel to the groups and final extended project team. Arrows indicate these communication iterations and information transference between facets of the project.
